# Impact of opioid-based versus opioid-free anaesthesia on postoperative recovery, circulating inflammatory cytokines and brain-derived neurotrophic factor

**DOI:** 10.64898/2026.09.25.754530

**Authors:** Yann Gricourt, Kenneth Lin, Thorsten Althoff, Eileen Nguyen, Andrea Mattern, Meenakshi Nair, Tess Leong, Emily Silva, Tristan Grogan, Nicolas Massaly

**Affiliations:** Department of Anaesthesiology & Perioperative Medicine, David Geffen School of Medicine, University of California Los Angeles, Los Angeles, USA; Department of Anaesthesiology and perioperative medicine, Nimes University Hospital, Nimes, France; IMAGINE UR UM 103, Montpellier University, Anaesthesia Critical Care, Emergency and Pain Medicine Division, Nimes University Hospital, Nimes, France; Division of Laboratory Animal Medicine, University of California Los Angeles, Los Angeles, USA; Department of Medicine Statistics Core, David Geffen School of Medicine, University of California Los Angeles, Los Angeles, USA

**Keywords:** Opioid-Free Anaesthesia, Opioid-Based Anaesthesia, Dexmedetomidine, BDNF, Quality of Recovery

## Abstract

**Background:** Opioid-free anaesthesia (OFA) has emerged as an alternative to opioid-based anaesthesia (OBA), but its benefit over OBA for postoperative quality of recovery (QoR) remains unclear. To address this question while limiting variability inherent in clinical studies, such as analgesic regimens and patient psychosocial factors, we investigated the effects of OFA versus OBA on postoperative recovery in a controlled preclinical mouse model.

**Methods:** Adult male and female C57BL/6J mice underwent laparotomy or sham procedures under isoflurane anaesthesia, and were randomly assigned to saline (control), fentanyl (OBA), or dexmedetomidine (OFA) as intraoperative analgesic adjuncts. All mice received multimodal postoperative analgesia, including carprofen. We adapted a standardised multidimensional recovery model using six behavioural endpoints reflecting human QoR domains: body weight, food intake, locomotor activity, mechanical sensitivity, sucrose preference, and social interaction. These endpoints were assessed on postoperative days 1, 2, 3, and 7. Plasma inflammatory cytokine and brain-derived neurotrophic factor (BDNF) concentrations were also measured in parallel as potential biological correlates of recovery.

**Results:** Over the study period, we observed no treatment effect across behavioural endpoints in laparotomised mice, including body weight (P=0.157), food intake (P=0.884), locomotor activity (P=0.864), mechanical sensitivity (P=0.628), sucrose preference (P=0.615), and social interaction (P=0.442). Composite QoR scores did not differ between OFA and OBA groups (P=0.569), with no differences in postoperative circulating inflammatory cytokine or BDNF concentrations.

**Conclusions:** Dexmedetomidine-based OFA and fentanyl-based OBA produced similar postoperative behavioural recovery, circulating inflammatory cytokine and BDNF trajectories in laparotomised mice receiving multimodal postoperative analgesia.

## INTRODUCTION

While opioids remain the standard analgesic adjuvant for intraoperative nociception control, their adverse effects have prompted increasing interest in opioid-free anaesthesia (OFA) strategies, including regional anaesthesia and non-opioid analgesic adjuvants. ^1–5^ Among these, dexmedetomidine, a selective α2-adrenergic receptor agonist with both analgesic and sedative properties, has been associated with reduced perioperative inflammation and improved quality of recovery (QoR) in humans. ^6–8^ Nevertheless, whether dexmedetomidine-based OFA provides analgesia equivalent to standard opioid-based anaesthesia (OBA) and presents meaningful benefits beyond reducing postoperative nausea and vomiting remains actively debated. ^4, 9–11^ Clinical studies comparing postoperative QoR following OFA versus OBA have yielded conflicting results ^12, 13^ which may be reflecting differences in surgical procedures, analgesic regimens, patients’ psychosocial factors and the subjective nature of QoR assessments. ^14^

Here, we developed a multidimensional mouse model of postoperative recovery following laparotomy, adapted from a previously established rat model, ^15^ to address these limitations. With this model, we assessed multiple dimensions of postoperative recovery reflecting domains captured by the clinically used Quality of Recovery-15 score, ^16^ including physical activity, food intake, hyperalgesia, hedonic states, and social interaction.

The current study aims at investigating how intraoperative dexmedetomidine-based OFA or fentanyl-based OBA influences recovery trajectories. Importantly, a multimodal analgesia was provided to all animals for two days after laparotomy to accurately reverse translate the standard perioperative care in humans, and to allow us to isolate the contribution of the intraoperative analgesic choice on recovery outcomes.

Beyond behavioural recovery, circulating biological markers, such as cytokines, provide a complementary and objective measures of the physiological response to surgery. ^17^ Indeed, surgical procedures lead to inflammatory and neurobiological responses that can persist into the postoperative period. ^18–21^ Circulating inflammatory cytokines are reliable indicators of postoperative immune activation, ^17^ whereas levels of brain-derived neurotrophic factor (BDNF), a protein involved in neuroplasticity and cognitive processes, have been reported to decrease following surgical procedure. ^19, 20, 22^ However, the impact of OFA and OBA on these circulating markers during the postoperative recovery period remains untested.

The current study has two main objectives. The first aim is to determine whether dexmedetomidine-based OFA, compared to the standard intraoperative OBA regimen, alters postoperative recovery in mice undergoing laparotomy. The second aim is to uncover whether the OBA and OFA strategies differentially influence postoperative circulating inflammatory cytokine and BDNF trajectories.

Our findings demonstrate that, while using a standardized postoperative analgesia to mimic patient conditions, intraoperative OFA leads to postoperative recovery trajectories and circulating inflammatory cytokines and BDNF levels comparable to those observed using an OBA strategy. Overall, our results highlight dexmedetomidine-based intraoperative analgesia as a promising option for selective patients for whom opioids should be avoided without compromising their postoperative recovery.

## MATERIALS AND METHODS

### Animals

All procedures were approved by the University of California, Los Angeles Institutional Animal Care and Use Committee (IACUC) in accordance with the National Institutes of Health Guidelines for the Care and Use of Laboratory Animals under the protocol ARC-2022-088. The completed ARRIVE 2.0 guidelines checklist is provided in Supplementary Data 1.^23^ Adult male and female C57BL/6J mice, 8–12 weeks old with a minimum body weight of 18 g at the start of the experiments, were obtained from the Jackson Laboratory. Both sexes were included in all experiments. Because the study was not powered for sex-specific comparisons and sex was not a prespecified primary analytic factor, males and females were pooled for the primary analyses. Mice were group housed (4 animals per cage) on a 12/12-hour dark/light cycle (lights on at 7:00 AM). Mice had ad libitum access to food and water and were habituated to the vivarium for at least one week before the start of the experiments.

Five cohorts were included, each lasting 2 weeks (habituation and postintervention assessment). A total of 64 mice were included in the analysis. Mice presenting complications during surgical procedures or blood collection were excluded from analysis.

Treatments and groups were randomly assigned to all mice using computer-generated randomisation by an independent investigator. Surgical procedures and blood collections were each performed by the same experimenter, while behavioural assessments were consistently performed by the same two experimenters. All experimenters and statistician remained blinded to treatment allocation until completion of data analysis, to the best of our abilities as sham and control mice did not have ventral abdominal sutures.

### Anaesthesia and surgical procedures

On the surgical day (postoperative day 0: POD0), mice were allocated to the three predefined groups: control, sham surgery, and surgery.

Control mice received no anaesthesia and no surgery but ran all behavioural assessments and blood analyses. The sham group received preoperative care, anaesthesia but no surgical procedure, and ran both behavioural assessments and blood analysis. Lastly, the surgery group underwent similar experimental procedures and laparotomy. All mice received multimodal postoperative analgesic regimen, consisting of intraperitoneal bupivacaine (2 mg.kg ¹) and nonsteroidal anti-inflammatory drug (carprofen, 5 mg.kg ¹) 15 minutes before surgical procedure, on Post Operative Day (POD) 1 and 2 to model pre and postoperative patient care and isolate the impact of intraoperative OBA and OFA during recovery.

Both sham and surgery groups received either 0.9% sterile saline, Dex (25 µg.kg ¹; OFA group), or fentanyl (0.2 mg.kg ¹; OBA group), administered at a volume of 10 mL.kg ¹, immediately after anaesthesia induction. Given its reported benefits on postoperative recovery and circulating BDNF levels, Dex was selected as the non-opioid adjuvant for this study. ^7, 24^. Anaesthesia was initiated using 2% isoflurane mixed with 95% O_2_ and 5% CO_2_ (2 L.min ¹) and lasted for 45 minutes. For surgical and sham procedures, mice were placed supine on a heating pad, abdominal fur was then removed, and the skin was prepared using an antiseptic solution. Anaesthetic depth was assessed by loss of the paw withdrawal reflex. Heart rate (HR), Breath rate (BR), and arterial oxygen saturation (O_2_Sat) were continuously monitored throughout the procedure using a non-invasive pulse oximeter (Starr Life Sciences, USA). ^25–27^ Mice allocated to the surgery group underwent a standardised laparotomy procedure, ^28^ a validated model for investigating postoperative pain mechanisms.^29^

Briefly, for laparotomy procedure mice received a ∼2-cm incision of the midline abdominal skin and muscle layers 20 minutes after anaesthesia induction and treatment administration, when mice were deeply sedated. A gentle manipulation of the bowel was performed for 30 seconds using sterile haemostats. Muscle and skin layers were sutured back together using sterile absorbable sutures. Mice allocated to Sham condition did not undergo laparotomy and were used to assess the effects of OBA vs OFA treatments on behavioural outcomes, circulating cytokines and BDNF, independent of surgical procedure.

After 45 min of sedation, all mice (sham and surgery groups) were placed on their back in a clean recovery cage placed on top of a water circulating blanket. To assess the impact of OBA and OFA on anaesthesia emergence, latency for return of righting reflex (RORR) and first ambulatory movement ^30^ were video recorded and manually scored.

### Blood Collection and cytokines analysis

To assess postoperative circulating inflammatory biomarkers and BDNF levels, 60 µL of whole blood was collected right before the surgical procedure (POD0) (excluding control mice) and on POD 1, 2, 3, and 7 in all mice. Blood samples were collected using 100µl calibrated pipets (VWR international, cat no 53432-921) containing 5µl of Ultrapure 0.5M EDTA, pH 8.0 (Invitrogen, cat no 15575-038). Retro-orbital sinus (under anaesthesia), submental and facial vein punctures (restrain), were used to collect blood on POD0, 1, 2, and 3, respectively.^31^ Lastly, blood was collected at POD7 using the contralateral retro-orbital sinus, under brief non-survival anaesthesia. Each blood collection site was used only once. All blood samples were collected at the same time each day to avoid circadian impact on circulating biomarkers.

After collection, blood samples were kept on ice and centrifuged at 4 °C for 10 minutes at 2,000×g. The pellet was discarded, and the supernatant was centrifuged again at 2,000×g. After this second centrifugation, the supernatant was stored at −80 °C until analysis. ^32^ Circulating levels of cytokines and chemokines were determined using an in-house 32-Plex mouse cytokine/chemokine ELISA kit (assessing Eotaxin/CCL11, G-CSF, GM-CSF, IFN-γ, IL-1α, IL-1β, IL-2, IL-3, IL-4, IL-5, IL-6, IL-7, IL-9, IL-10, IL-12 (p40), IL-12 (p70), IL-13, IL-15, IL-17, IP-10, KC, LIF, LIX, MCP-1, M-CSF, MIG, MIP-1α, MIP-1β, MIP-2, RANTES, TNF-α, VEGF) from the Immunogenetics Core at the University of California, Los Angeles. BDNF levels were determined using Mature BDNF ELISA kits (Fujifilm, Cat. #290-85801). All tests were run per manufacturer recommendations.

### Behavioural assays

Behavioural procedures were designed to evaluate postoperative recovery, as previously reported in rats, ^15^ and to reflect domains evaluated by the Quality of Recovery-15 score in humans, including physical comfort, physical independence, pain, emotional state, and psychological support. ^16^ While some human recovery domains cannot be directly modelled in rodents, such as psychological support from healthcare providers, the behavioural battery was designed to capture selected aspects of physical and emotional recovery. Von Frey testing, locomotor activity, food consumption, and body weight were used to assess physical recovery, whereas sucrose preference and social interaction were used to explore affective states during recovery.^16^

For all behavioural procedures, three daily habituation sessions and baseline data collected (fourth session) were performed before POD0. The exact sequence of behavioural procedures was repeated on POD1, 2, 3, and 7 (see **Fig. 1**). All mice returned to their home cages and vivarium daily after test completion. Body weight was recorded twice daily (morning and evening) throughout the study.

**Fig. 1.**
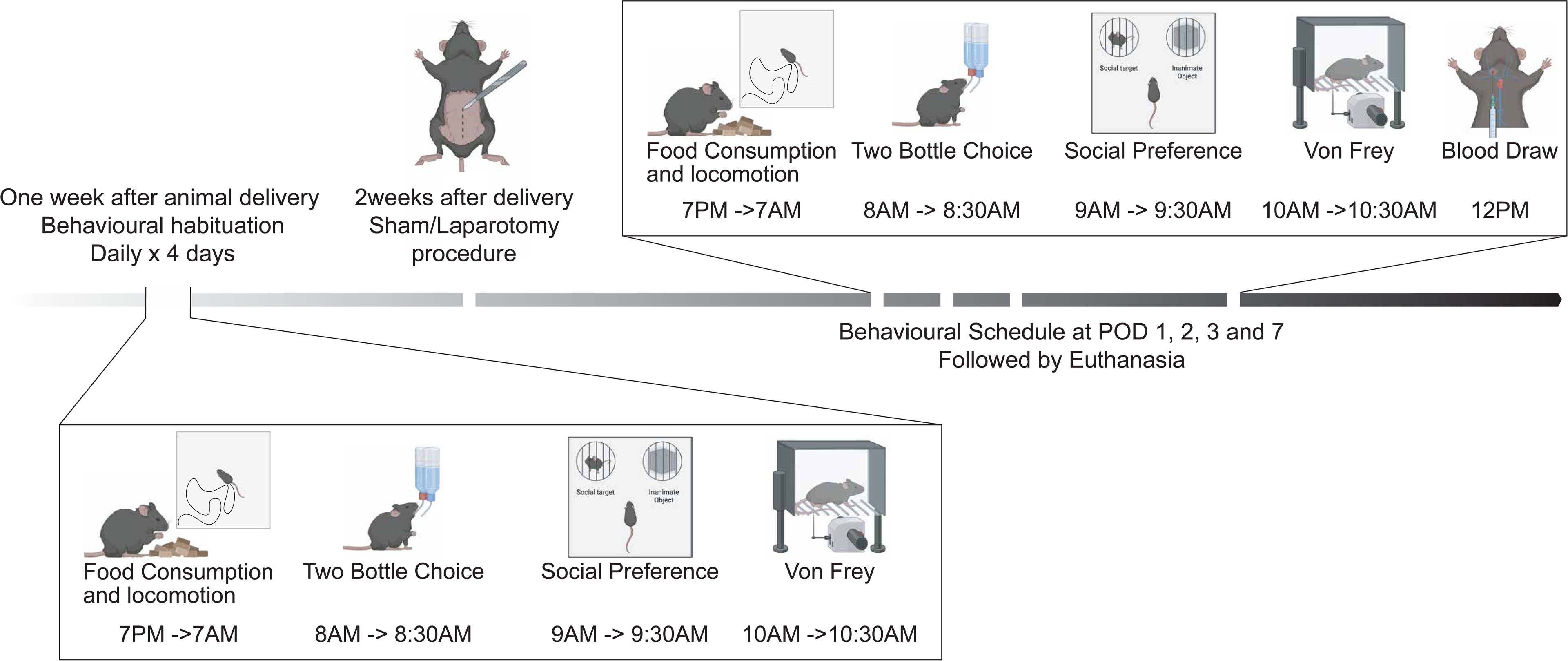
Experimental timeline and postoperative behavioural assessment procedures. Mice underwent four daily habituation sessions before surgery. Behavioural assessments were then performed on Post Operative Day (POD) 1, POD2, POD3, and POD7 following a standardized sequence including overnight food consumption and locomotor activity, body weight, two-bottle choice, social preference, Von Frey testing, and blood sampling. Mice were euthanized on POD7.

### A- Food self-administration

To monitor food self-administration as a marker of physical comfort, mice were single-housed overnight (7:00 PM–7:00 AM) in a home cage–like setting equipped with an operant feeder - FED3 device - affixed to the cage’s wall. ^33^ The FED3 device was equipped with two nose-poke ports and a food receptacle. Under a Fixed-Ratio 1 (FR1) schedule of reinforcement, interaction with the active port resulted in delivery of a food pellet (20-mg sucrose pellet; Bio-Serv) in the food receptacle, whereas interaction with the inactive port had no consequence. Subsequent rewards were only delivered after retrieval of pellets from the food receptacle to accurately monitor food consumption. Following the overnight session, mice were placed back in their home-cage.

### B – Locomotor activity

Locomotor activity (total distance travelled) was recorded and analysed using ANY-maze software during overnight session (mice’s active phase) and expressed as a percentage of baseline locomotor activity, accounting for intra-individual performances.

### C - Two-Bottle Choice

A two-bottle sucrose preference test (2BC) was used as a behavioural proxy for anhedonia. ^34^ To stimulate liquid seeking and consumption mice were water-restricted during the overnight food self-administration session, as previously reported. ^34^ An hour after completion of the food self-administration session, mice were individually placed in a home cage–like setting with bedding and given free access to either water or a 2% sucrose solution for 30 minutes using automated cage sippers. ^35^ Consumption was determined by the change in bottle weight (ΔW = weight before − weight after the session) for each solution. Sucrose preference was calculated as follows: (ΔW_sucrose / (ΔW_sucrose + ΔW_water)) × 100.

### D – Social Interaction

Following 2BC, mice were placed for 20 minutes in an open rectangular Plexiglas arena (50 × 30 × 23 cm) containing both a nonfamiliar, age-matched conspecific mouse placed under a wired enclosure, and a colour-matched object placed under a second wired enclosure, as a control. During the test, mice interactions with conspecific and object were monitored. Interaction was considered positive when the head of the test mouse was within 2 cm of either enclosure. Sociability index was calculated as: ((time exploring matched social partner)/(time exploring matched social partner + time exploring 3D control object)). ^36^

### E – Mechanical thresholds - Von Frey

After social interaction monitoring, paw withdrawal thresholds (PWTs) were measured using an electronic Von Frey (Ugo Basile) to assess for mechanical hyperalgesia.^37^ Mice were placed in individual Plexiglas chambers for 15 minutes, and mechanical sensitivity was assessed in quadruplicate on the right hind paw. PWTs were calculated as the mean value of the four measurements obtained during each testing session.

### F – Quality of recovery score

To reflect clinical recovery assessment, a composite QoR score was established based on a previously described rat model. ^15^ A pilot cohort exposed to similar conditions was used to define endpoint-specific maximum deficits as the lowest postoperative values (on POD1–2 for sucrose preference, overnight activity, and food intake; on POD2–3 for body weight, Von Frey threshold, and social interaction). Baseline values were obtained during the fourth habituation session. Outcomes were expressed as percentage of maximal possible effect (%MPE): %MPE = ((Test − baseline) / (maximum deficit − baseline)) × 100. Each endpoint was categorized into a 4-level ordinal score based on %MPE, with higher %MPE values indicating greater impairment: 0 (75–100%), 1 (50–74%), 2 (25–49%), and 3 (0–24%). To account for missing outcomes due to technical issues, endpoint scores were summed and divided by the number of available behavioural assessments per day, yielding a daily composite Quality of Recovery (QoR) score ranging from 0 to 3, with higher values indicating better recovery.

### Statistical analysis

All statistical analyses were performed by a blinded experimenter. Data analysis for intraoperative monitoring and emergence from sedation (**Fig. 2**) was performed using GraphPad Prism. Normal distribution of experimental data samples was assessed using the Shapiro–Wilk normality test. Datasets were then analysed with parametric or non-parametric comparisons, accordingly. Alterations in HR, BR, and O_2_Sat were assessed using a Two-way ANOVA. Changes in emergence from sedation were assessed using a One Way ANOVA. When ANOVA yielded a significant main effect or interaction, post-hoc comparisons were conducted.

**Fig. 2.**
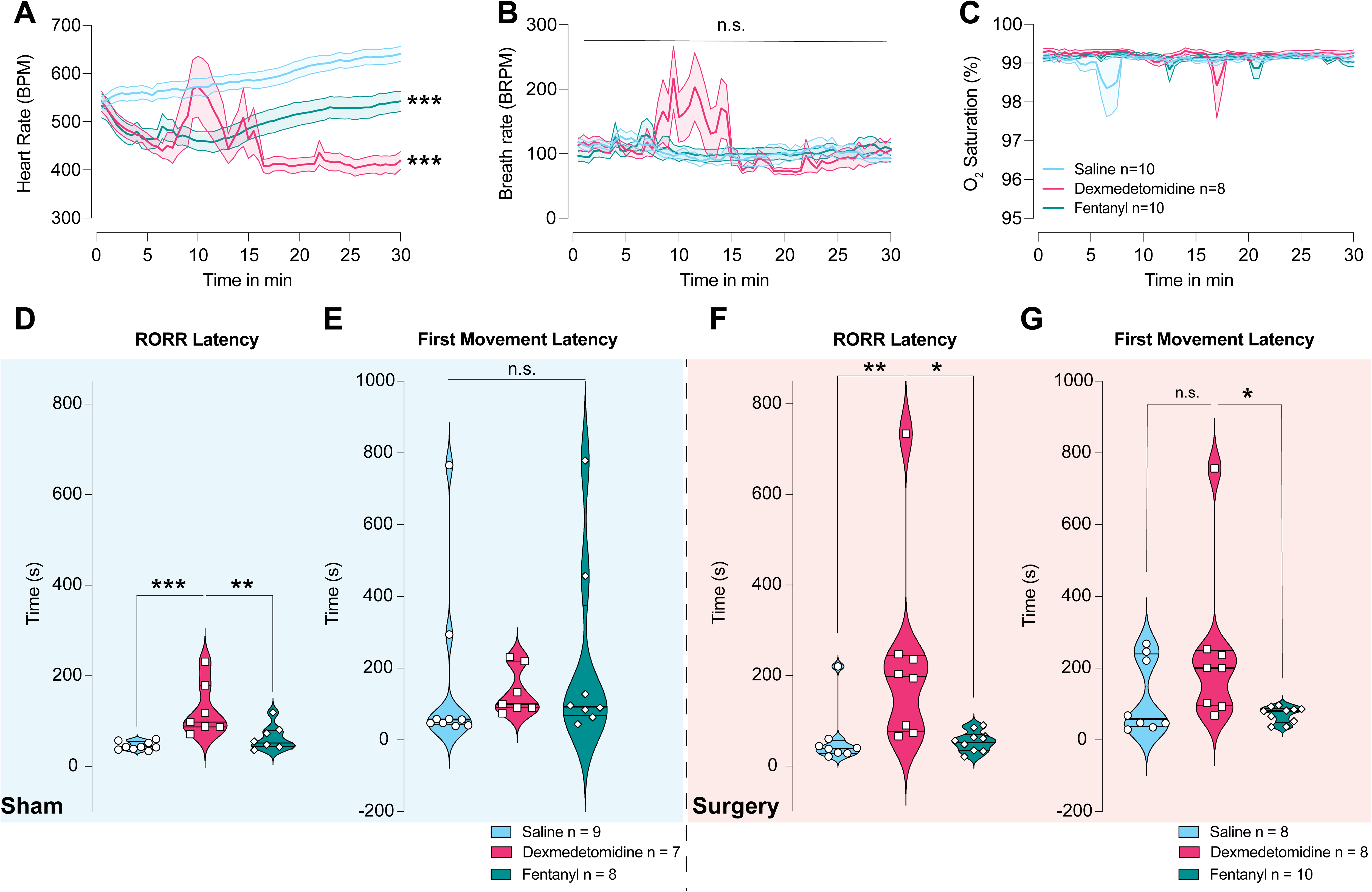
Dexmedetomidine induces bradycardia and delays emergence from anaesthesia. (A) Mean values of HR monitoring after OFA/OBA/control treatments. Both fentanyl (OBA) and Dex (OFA) decreased HR during the surgical procedure. B) Mean values of BR monitoring after OFA/OBA/control treatments. Anaesthetic strategy (OBA vs OFA vs control) did not impact the BR during the surgical procedure. C) Mean values of O_2_Sat monitoring after OFA, OBA, or control treatments. Anaesthetic strategy (OBA vs OFA vs control) did not impact the O2Sat during the surgical procedure. D) Dex (OFA, n=7) increased time until RORR compared to control (n=9) and fentanyl (OBA, n=8) groups during sham conditions. E) Anaesthetic strategy (OBA vs OFA vs control) did not impact the latency to first movement after anaesthesia in sham groups. F) Dex (OFA, n=8) increased RORR compared to control (n=8) and fentanyl (OBA, n=10) groups after the surgical procedure. G) Dex (OFA, n=8) increased the latency to first movement after the surgical procedure compared to fentanyl (OBA, n=10) group. All data are represented as mean values +/- S.E.M. * p<0.05; ** p<0.01; *** p<0.001; n.s. no significant difference p>0.05. Legend: BR: breath rate; HR: heart rate; O2Sat: oxygen saturation; RORR: return of righting reflex

Longitudinal behavioural and cytokine outcomes were analysed in R version 4.4.3 using linear mixed-effects models with a random intercept for mouse to account for repeated measurements over time. For the primary treatment analyses, untreated control mice were excluded and models included fixed effects for surgery group, treatment group (saline, Dex, fentanyl), time, and all interactions. Additional within-sham and within-surgery mixed-effects models including treatment, time, and treatment-by-time interaction terms were used to assess whether treatment trajectories differed over time within each surgical condition. Complementary control-inclusive mixed-effects models using a combined group variable were used for contextual comparisons and figure generation. Post hoc pairwise contrasts were obtained from estimated marginal means with Holm adjustment when appropriate. Cytokine outcomes were analysed on the log(1+x) scale to reduce skewness and better satisfy the statistical assumptions of the models. All tests were two-sided and P < 0.05 was considered statistically significant.

Sample sizes were selected based on prior postoperative behavioural studies in rodents,^15^ feasibility of repeated longitudinal behavioural testing and serial blood sampling, and available laboratory resources. Group sizes for the behavioural recovery experiments were generally 7–11 mice per group. As a rough benchmark, a sample size of 10 mice per group provides approximately 80% power to detect a standardized effect size of about 1.3 using a two-sample t-test with a two-sided alpha of 0.05. This is a conservative simplification of the longitudinal mixed-effects models used in the present study, which leverage repeated measurements over time. Therefore, the study was most sensitive to relatively large group differences, and smaller effects may not have been detectable.

## RESULTS

### Dexmedetomidine induces bradycardia and delays emergence from anaesthesia

First, to evaluate intraoperative safety and hemodynamic profiles associated with OFA versus OBA we monitored Heart Rate (HR), Breath Rate (BR), and Oxygen Saturation (O_2_Sat) during anaesthesia maintenance. This is critical as Dex is a selective α2-adrenergic agonist, well documented to induce dose-dependent bradycardia. ^7, 38, 39^

HR was significantly decreased after both fentanyl and Dex treatments compared to saline (two-way ANOVA: treatment effect, F_(2,25)_ = 13.82, p < 0.001; treatment × time interaction, F_(6.19,77.36)_ = 8.15, p < 0.001) (**Fig. 2A**). In contrast, BR and O Sat were not impacted by either treatment (BR: F_(2,25)_ = 0.31, p = 0.735; O_2_Sat: F_(2,26)_ = 0.94, p = 0.405; **Fig. 2B-C**).

To characterize the impact of OFA and OBA on anaesthesia emergence, we monitored latency to Return Of Righting Reflex (RORR) and first ambulatory movement. Dex significantly prolonged latency to RORR in both sham (one-way ANOVA: F_(2,21)_ = 10.44, p = 0.0007; Tukey: saline vs Dex, p = 0.0007; Dex vs fentanyl, p = 0.0079) and surgery groups (Kruskal–Wallis p = 0.0018; Dunn: saline vs Dex, p = 0.0026; Dex vs fentanyl, p = 0.0159) (**Fig. 2D and F, respectively**). In contrast, latency to first ambulatory movement was not significantly altered in sham mice (Kruskal–Wallis, p = 0.101) and showed only a modest difference after surgery (Kruskal–Wallis, p = 0.036) (**Fig. 2E and G, respectively**). This effect was driven by a significant increase in first movement observed after OFA (Dex) compared to OBA (fentanyl) (p = 0.046), while no difference was observed between OFA (Dex) and saline treatments (p = 0.133).

### Intraoperative dexmedetomidine does not improve postoperative recovery compared with fentanyl

The benefits of OFA versus OBA on postoperative QoR remain debated in clinical studies. ^4, 10, 11^ To address this question using a preclinical clinical, we investigated longitudinal postoperative recovery using multiple evoked and non-evoked behavioural endpoints and a composite QoR score.

#### A - Body weight, overnight food intake and locomotor activity

Laparotomy significantly decreased mice’ body weight (p<0.001) and food intake (p<0.001) during the early postoperative period, compared with sham controls (**Fig. 3A-B**), consistent with previous reports. ^15, 40^ These values returned to levels comparable to baseline around POD7.

**Fig. 3.**
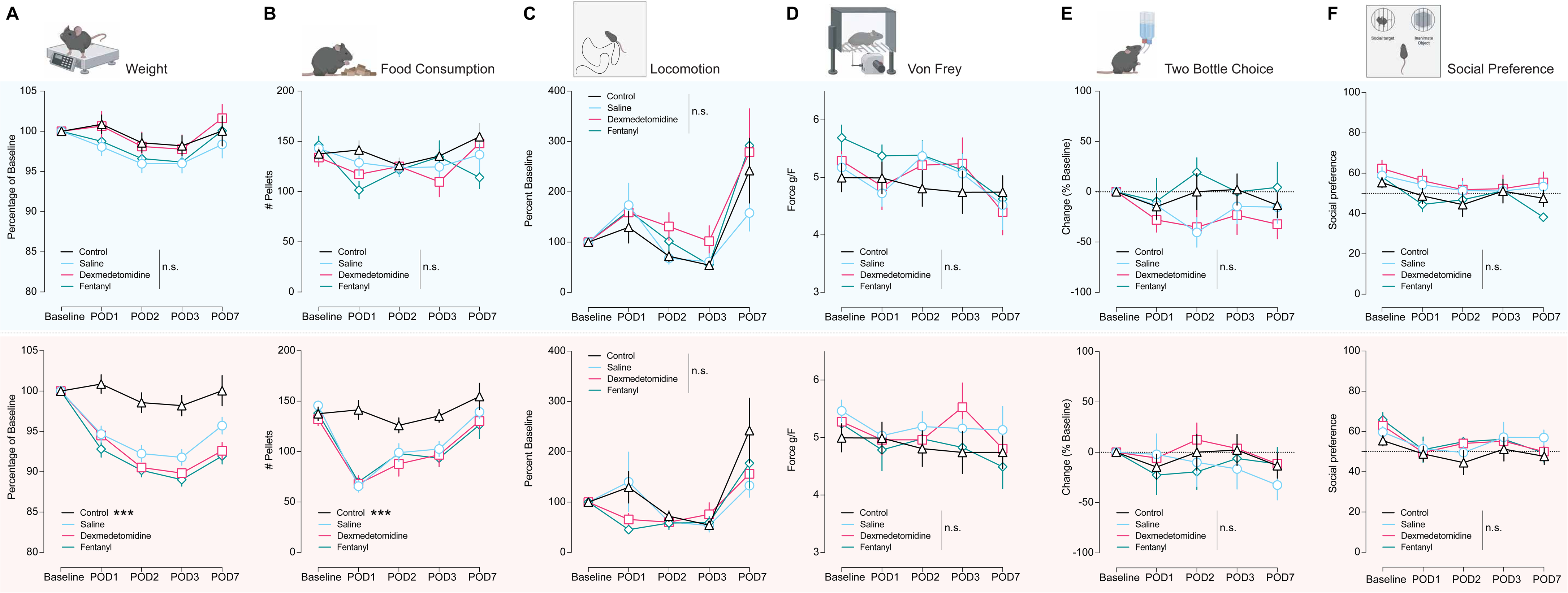
Intraoperative anaesthetic strategy does not impact mouse weight, food consumption, locomotor activity, mechanical threshold, and anhedonia after anaesthesia. For all graphical representations, sham groups are presented in top panels (blue), and surgery-exposed groups are presented in bottom panels (red). Baseline and postoperative assessments were performed before anaesthesia and on POD1, POD2, POD3, and POD7, respectively. A) Percent of baseline weight. B) Number of food pellets (20 mg per pellet) consumed during 12-hour overnight sessions. C) Percent of baseline horizontal locomotor activity. D) Plantar mechanical thresholds measured using Von Frey testing. E) Percent change in sucrose preference over water. F) Percent social preference over inanimate object. All data are represented as mean values ± S.E.M. n.s. no significant difference p>0.05. Legend: POD: postoperative day

The intraoperative treatment (OFA vs OBA vs Saline) administered before laparotomy did not impact surgery-induced body weight (mixed-effects model for repeated measures, drug x time interaction effect: p = 0.157) or food intake (mixed-effects model for repeated measures, drug x time interaction effect: p = 0.884) decrease (**Fig. 3A–B, bottom**). Similarly, in sham mice, neither Dex nor fentanyl significantly altered body weight (mixed-effects model for repeated measures, drug x time interaction effect: p = 0.712) or food intake (mixed-effects model for repeated measures, drug x time interaction effect: p = 0.171) compared with saline treated animals (**Fig. 3A–B, top**).

In addition to body weight and food intake, postoperative pain is associated with decreased ambulatory activity in rodents, an alteration that can be prevented with adequate analgesic management. ^41^ Here we demonstrate that neither intraoperative OFA nor OBA altered locomotion in mice who underwent laparotomy (mixed-effects model for repeated measures, drug x time interaction effect: p = 0.864) or in control (sham) animals (mixed-effects model for repeated measures, drug x time interaction effect: p = 0.101) (**Fig. 3C**).

#### B - Mechanical sensitivity and affective measures

To assess whether the intraoperative anaesthesia regimen impacted laparotomy-induced mechanical hyperalgesia, mice were exposed to plantar Von Frey stimulation. Our data showed that laparotomy did not significantly alter mechanical thresholds (**Fig. 3D, bottom)**, and neither Saline, OFA, nor OBA altered mechanical sensitivity in either sham or laparotomy groups (Sham: mixed-effects model for repeated measures, drug x time interaction effect: p = 0.899; Surgery: mixed-effects model for repeated measures, drug x time interaction effect: p = 0.628; **Fig. 3D**).

To investigate whether intraoperative treatment could impact post-surgical anhedonia, defined as a decrease in interest in a previously rewarding activity, mice were exposed to two-bottle choice (2BC) and social interaction tests, two well-validated preclinical models for assessing anhedonic states. ^34, 42^ Overall, neither sucrose preference nor social interaction with a conspecific was impacted by laparotomy procedure (**Fig. 3E-F, bottom**). Further, neither Saline, OBA, nor OFA impacted these behavioural outcomes in either sham (mixed-effects model for repeated measures, drug x time interaction effect: for sucrose preference: p = 0.534; for social interaction: p = 0.481) or laparotomy exposed mice (mixed-effects model for repeated measures, drug x time interaction effect: for sucrose preference: p = 0.615; for social interaction: p = 0.442) (**Fig.3 E-F**).

#### C - Total QoR score

To reflect the multidimensional clinical recovery assessment, we established a composite recovery score based on the above described behavioural and sensory measures, each contributing equally to the daily composite QoR score, as shown previously in rats. ^15^ In control-inclusive longitudinal mixed-effects models, mice exposed to laparotomy showed worse early recovery trajectories than untreated controls (group × time p = 0.023) (**Fig. 4B**). However, no significant intraoperative treatment effect was observed on QoR scores within sham mice (mixed-effects model for repeated measures, drug × time p = 0.227) or laparotomy exposed mice (mixed-effects model for repeated measures, drug × time p = 0.569) (**Fig. 4A-B).**

**Fig. 4.**
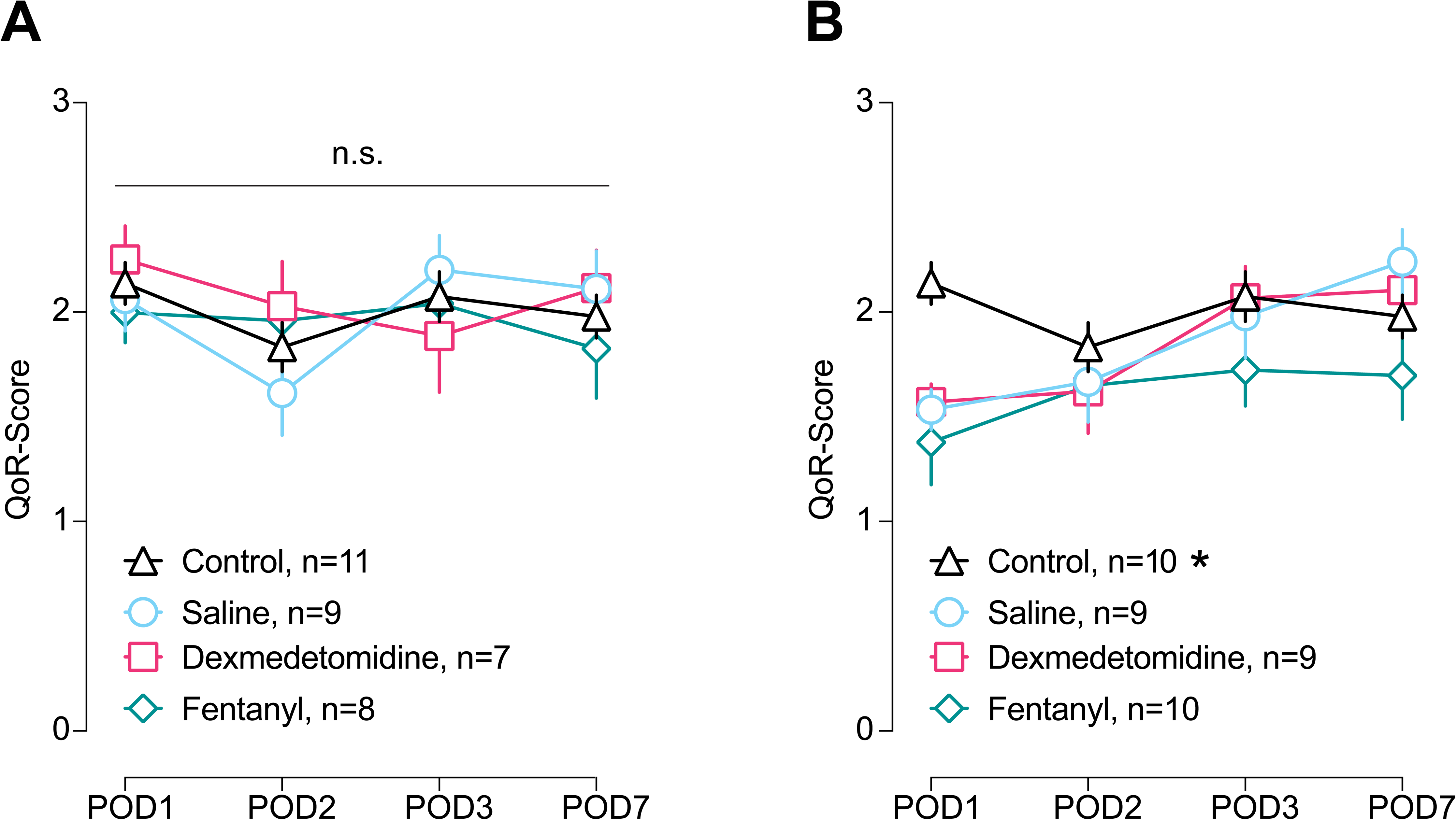
Intraoperative anaesthetic strategy does not impact overall post-operative recovery. Intraoperative OFA vs OBA vs control did not impact composite QoR score in mice undergoing either A) sham or B) surgery. All data are represented as mean values ± S.E.M. n.s. no significant difference p>0.05. Legend: OBA: opioid-based anaesthesia; OFA: opioid-free anaesthesia; QoR: quality of recovery

### Intraoperative opioid strategy does not alter circulating inflammatory cytokine levels

We next assessed whether intraoperative OBA and OFA treatments impacted circulating inflammatory markers after laparotomy. Importantly, no differences were observed across treatments for interferon-γ (mixed-effects model for repeated measures, drug x time interaction effect: p = 0.930), interleukin-1α (mixed-effects model for repeated measures, drug x time interaction effect: p = 0.154), interleukin-6 (mixed-effects model for repeated measures, drug x time interaction effect: p = 0.817), or TNF-α (mixed-effects model for repeated measures, drug x time interaction effect: p = 0.825) in sham animals (**Fig. 5A–D – Top**).

**Fig. 5.**
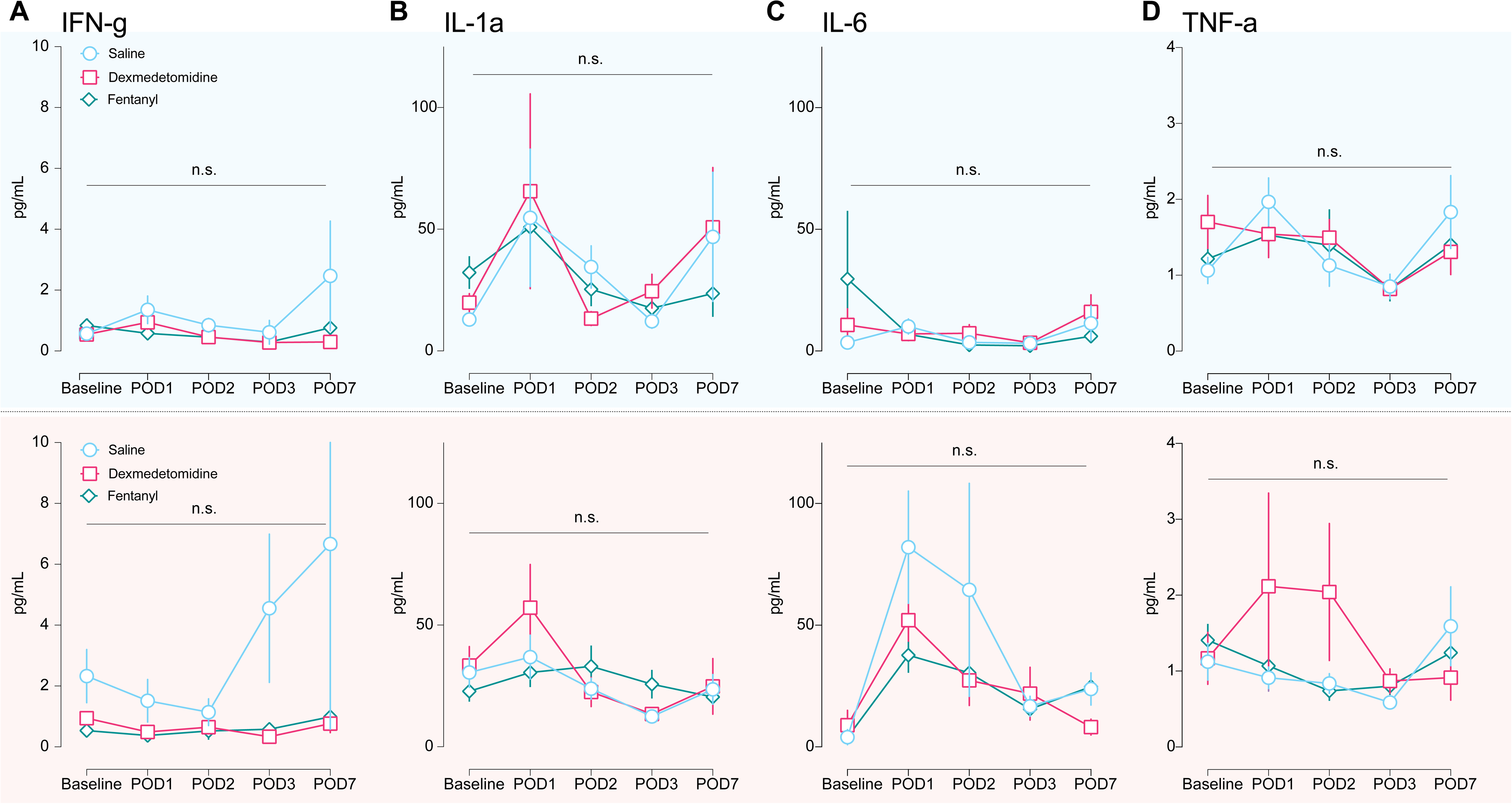
Intraoperative anaesthetic strategy does not impact circulating inflammatory markers. For all graphical representations, sham groups are presented in top panels (blue), and surgery-exposed groups are presented in bottom panels (red). Intraoperative OFA vs OBA vs control did not change circulating levels of A) IFN-γ, B) IL-1α, C) IL-6, and D) TNF-α on POD1, POD2, POD3, and POD7. All data are represented as mean values ± S.E.M. n.s. no significant difference p > 0.05. Legend: IFN-γ: interferon-gamma; IL-1α: interleukin 1-alpha; IL-6: interleukin 6; OBA: opioid-based anaesthesia; OFA: opioid-free anaesthesia; POD: postoperative day; TNF-α: tumour necrosis factor alpha.

Similarly, neither Saline, OBA, nor OFA impacted interferon-γ (mixed-effects model for repeated measures, drug x time interaction effect: p = 0.934), interleukin-1α (mixed-effects model for repeated measures, drug x time interaction effect: p = 0.315), and TNF-α (mixed-effects model for repeated measures, drug x time interaction effect: p = 0.973) levels in laparotomy exposed animals (**Fig. 5A–D – Bottom**).

Interestingly, while circulating IL-6 levels increased after laparotomy compared to baseline, neither Saline, Dex, nor fentanyl intraoperative treatment impacted these laparotomy-induced changes (mixed-effects model for repeated measures: Drug p = 0.519; Time p < 0.001; Drug × Time p = 0.904). Across treatments, circulating levels of IL-6 were higher at POD1 (estimate 2.80, p < 0.0001), POD2 (estimate 2.01, p < 0.0001), POD3 (estimate 1.64, p < 0.0001), and POD7 (estimate 0.86, p = 0.0021) than baseline levels.

### Intraoperative opioid strategy does not alter circulating BDNF levels

While reports demonstrated that BDNF levels decrease following surgery, a measure that has been correlated with poor perioperative outcomes such as delirium ^20^, whether intraoperative opioid strategy influences circulating BDNF levels remains unclear.^24^

Here, we demonstrate that neither Saline, OBA, nor OFA intraoperative treatments altered circulating BDNF levels in either sham (mixed-effects model for repeated measures, drug x time interaction effect: p = 0.507) or laparotomy-exposed mice (mixed-effects model for repeated measures, drug x time interaction effect: p = 0.927) (**Fig. 6A-B**). Interestingly and in contradiction with current literature, our data reveal a transient increase in circulating BDNF (POD1–3) in both sham and surgery groups, before returning to baseline levels at POD7.

**Fig. 6.**
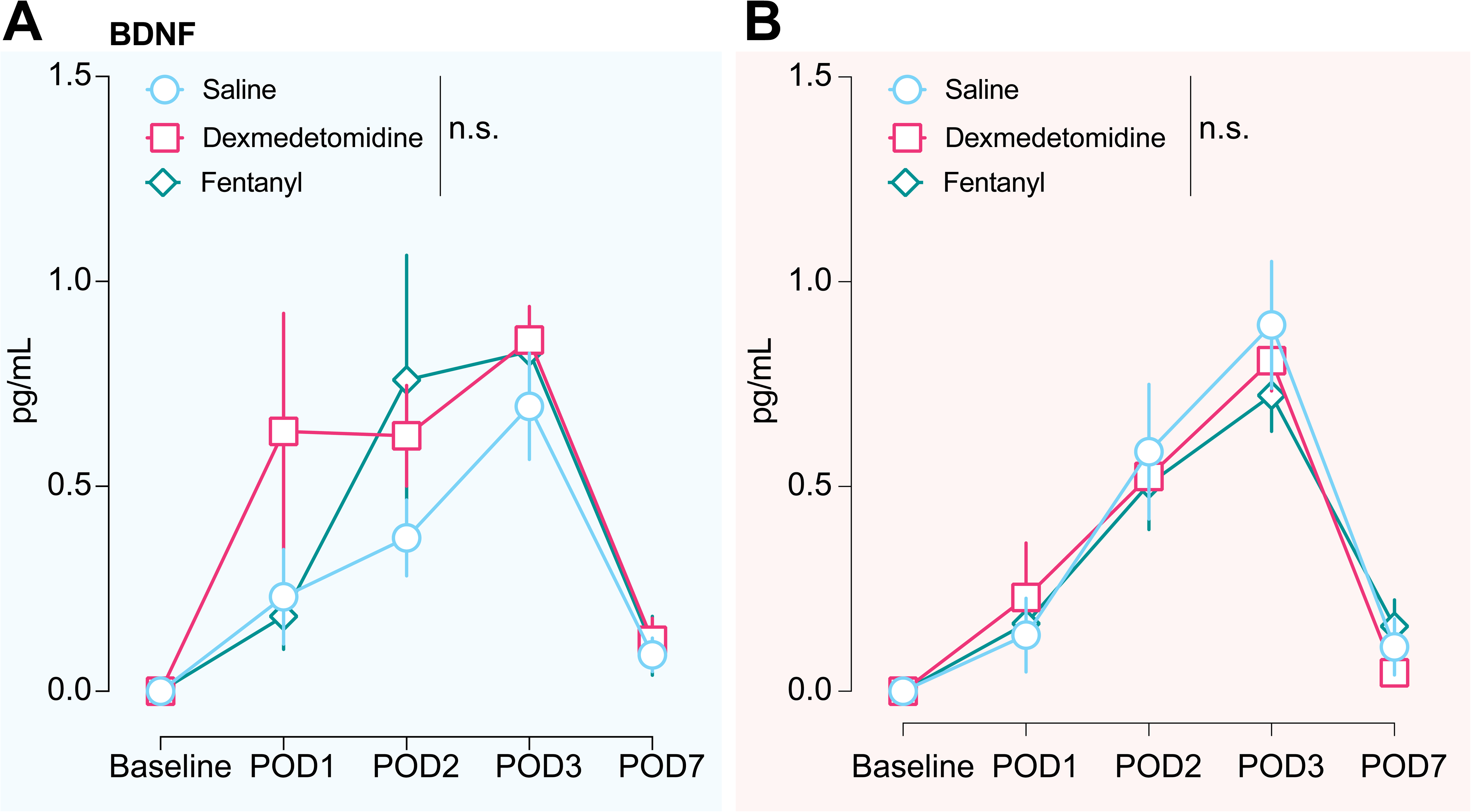
Intraoperative anaesthetic strategy does not impact circulating Brain-Derived Neurotrophic Factor level. Intraoperative OFA vs OBA vs control did not change circulating levels of BDNF in mice undergoing A) sham conditions, or B) surgery. All data are represented as mean values ± S.E.M. n.s. no significant difference p>0.05. Legend: BDNF: brain-derived neurotrophic factor; OBA: opioid-based anaesthesia; OFA: opioid-free anaesthesia

## DISCUSSION

In this study, we compared Dex-based (OFA) to fentanyl-based (OBA) anaesthesia regimens, the current standard of care in surgical practice, in post-laparotomy recovery trajectories and measured circulating inflammatory cytokine and BDNF. To do so, we developed a standardized behavioural model to investigate both sensory and affective components of postoperative pain in mice. Our findings demonstrate that OFA and OBA have similar recovery trajectories across multiple behavioural domains and overall composite score, consistent with clinical studies reporting limited or inconsistent clinical benefits associated with OFA. ^12, 13^ In addition, postsurgical circulating cytokines or BDNF trajectories do not differ between OFA and OBA. These results suggest that intraoperative OFA and OBA possess similar efficacy in laparotomy recovery when combined with appropriate postoperative analgesia.

### Dexmedetomidine-based OFA and fentanyl-based OBA have comparable postoperative recovery trajectories in mice

The results obtained in the early postoperative recovery period after laparotomy were consistent with other reports uncovering reduced body weight and no significant changes in hind paw mechanical sensitivity, sucrose preference, or locomotor activity. ^40, 43^ Several factors should still be considered regarding the lack differences observed on postoperative recovery in our settings **(Fig. 3-4)**. First, all laparotomy-exposed mice received standardized postoperative multimodal analgesia combining carprofen (5mg.kg^-1^) and bupivacaine (2mg.kg^-1^) for a couple of days. This postoperative treatment was included to improve translational relevance, as patients routinely receive analgesic care following such procedures. ^44^ However, this multimodal analgesic regimen may have attenuated potential differences between OFA and OBA in postoperative behavioural and circulating cytokine outcomes that could have emerged in the absence of postoperative analgesia. Second, although laparotomy is associated with acute postoperative pain, ^29, 45^ the surgical model used in this study may have induced limited nociceptive and inflammatory burden, as no organ resection or abrasion was performed. ^29^ Only body weight and food consumption were decreased following surgical procedure, while other behavioural outcomes were largely unaltered when compared to sham mice **(Fig. 3-4)**, limiting our ability to detect differences between anaesthetic strategies. A more invasive surgical model may reveal differences between OBA and OFA that were not apparent under the present conditions. Lastly, delayed recovery trajectories are more pronounced in older patients,^46^ warranting further investigation into whether Dex-based OFA may offer advantages over fentanyl-based OBA in an aged population. Such studies could provide important insights into optimizing anaesthetics strategies for older patients. Regarding safety-related outcomes, Dex-based OFA induced bradycardia and delayed anaesthesia emergence compared with other treatments **(Fig. 2)**, consistent with both preclinical and clinical studies. ^7, 8, 38, 39, 47^ Collectively, these findings emphasize the importance of considering and monitoring hemodynamic and emergence effects when both evaluating Dex-based OFA strategies in preclinical studies and using a Dex-based OFA strategy in clinical settings.

### Modelling multidimensional postoperative recovery in mice

To our knowledge, no previous study has developed a mouse model to systemically assess postoperative recovery following laparotomy or other surgical procedures. Adapted from a previously established rat model, ^15^ our approach was designed to investigate recovery dimensions commonly explored in clinical QoR, including physical, functional, and emotional aspects. ^16^ Other non-evoked behavioural procedures have been used to evaluate postoperative well-being in mice, including nesting behaviour, rearing activity, or wheel running. ^40^ These approaches showed no significant changes 24 h after laparotomy when compared with control mice, ^40^ limiting their utility for longitudinal assessment of postoperative recovery. Here we used additional automated, video-based behavioural approaches to capture distinct dimensions of postoperative recovery **(Fig. 3)** while reducing experimenter handling, which is known to impact pain-related behaviours in rodents. ^48^ The development of translationally relevant models of postoperative recovery is a critical objective given the growing importance of QoR assessments in perioperative settings. ^49^ While our model provides a framework for assessing multiple dimensions of recovery in mice, further validation across surgical procedures and clinically relevant interventions will be necessary to establish its translational relevance and utility.

### Postoperative circulating BDNF trajectories in mice

We demonstrate that circulating BDNF levels transiently increased during the early postoperative period in both sham and surgical mice, regardless of anaesthetic strategy, but remained below 1.0 pg.mL^-1^ throughout the study period (**Fig. 6**). This trajectory contrasts with postoperative BDNF decreases previously reported in humans. ^20^ Experimental stress and local tissue injury related to repeated blood sampling on submandibular and facial veins may partly explain this finding, as submandibular blood collection has been associated with increased circulating BDNF levels in stressed rats. ^50^ Further, the low BDNF concentrations obtained from mice blood samples in our study are consistent with the minimal BDNF content of mouse platelets. Indeed, in human patients, platelets are a major source of BDNF and substantially contribute to circulating levels of BDNF. ^32^ This striking difference may explain the inconsistency in postoperative circulating BDNF observed in mice and human patients.

In our hands, intraoperative Dex-based OFA did not affect postoperative circulating BDNF concentrations compared with fentanyl-based OBA (**Fig.6**). In sharp contrast, others have shown that Dex prevents surgery-induced decreases in BDNF levels in patients. ^24^ Similarly, Dex-based OFA has been shown to be sufficient to prevent BDNF decrease in the mice hippocampus after laparotomy. ^51^ As tissue and circulating BDNF levels may follow different trajectories in postoperative pain recovery, direct comparisons between our findings and this previous study remain limited, but warrant the need for better defining the relationship between peripheral and central BDNF and postoperative pain recovery.

In conclusion, intraoperative Dex-based OFA and fentanyl-based OBA produced similar postoperative behavioural recovery and circulating inflammatory cytokine and BDNF trajectories in laparotomised mice receiving multimodal postoperative analgesia. More broadly, our findings add critical evidence to support the use of intraoperative opioid-sparing strategies as a therapeutic option available to clinicians to tailor individualized anaesthetic care to each patient’s specific perioperative needs.

## Author contributions

- Conceptualization: YG, NM
- Methodology: YG, NM, KL
- Formal analysis: YG, KL, TG, NM
- Investigation: YG, KL, TA, AM, MN, TL, ES, NM
- Data curation: YG, KL, TA, NM
- Writing – Original Draft: YG, KL, NM
- Writing – Review & Editing: YG, KL, TA, EN, MN, TL, ES, NM
- Supervision: NM
- Project administration: NM

## Acknowledgments

We would like to thank all members of the Massaly lab and the Department of Anaesthesiology and Perioperative Medicine at University of California, Los Angeles for their helpful comments and discussions throughout the completion of the study. We would also like to thank the Immune Assessment Core at University of California, Los Angeles for their help in processing samples for inflammatory markers and BDNF.

## Conflicts of interest statement

YG receives speaker fees from GE Healthcare (Buc, France). The other co-authors have no conflicts of interests.

## Declaration of Generative AI and AI-assisted technologies in the writing process

During the preparation of this work the author(s) used Chat-GPT 5.5 in order to improve English readability. After using this tool/service, the author(s) reviewed and edited the content as needed and take(s) full responsibility for the content of the publication.

## Funding

This work was supported by the Department of Anaesthesiology and Perioperative Medicine and the University of California, Los Angeles start-up funds (NM), the University of Montpellier (YG; EXPLORE#6 grant), the Nimes University hospital (YG) and the Philippe Foundation (YG).

## Abbreviations

2BC: Two-Bottle Choice
BDNF: Brain-Derived Neurotrophic Factor
BR: Breath Rate
Dex: Dexmedetomidine
HR: Heart Rate
O2Sat: Oxygen Saturation
OBA: Opioid-Based Anaesthesia
OFA: Opioid-Free Anaesthesia
POD: Postoperative Day
PWT: Paw Withdrawal Threshold
QoR: Quality of Recovery
RORR: Return of Righting Reflex
RR: Respiratory Rate

